# Cardioprotective effects of AMPK activation in H1N1 influenza virus infection

**DOI:** 10.1101/2025.08.28.672931

**Authors:** Steve Leumi, Matthew McFadden, Naresh Kumar, Sameer Salam Mattoo, Jack Roettger, Samuel Speaks, Benjamin Liu, Shreenath Mohan, Brendan Reznik, Jana M. Cable, Parker J. Denz, Murugesan V. S. Rajaram, Adriana Forero, Jacob S. Yount

## Abstract

Cardiac complications are among the most common and severe extrapulmonary manifestations of influenza virus infection, yet they are rarely recapitulated in mouse models without immunodeficiency. We found that influenza virus A/California/04/2009 (H1N1) carrying a mouse-adaptive amino acid substitution in the PB2 protein (E158A) disseminates to the heart in WT C57BL/6 mice, where it induces inflammation, electrical dysfunction, and fibrotic remodeling. Influenza virus-infected heart tissue was significantly altered in mitochondrial metabolism, extracellular matrix, circadian rhythm, and immunity pathways. Particularly striking was activation of gene expression downstream of the mitochondrial biogenesis-promoting AMPK/PGC-1α axis, which occurred late in infection but failed to reverse the repression of mitochondria-associated genes, suggesting an insufficient or delayed compensatory response. Accordingly, we administered AMPK activator 5-aminoimidazole-4-carboxamide ribonucleoside (AICAR) early in infection and observed restoration of mitochondria-associated gene levels, amelioration of cardiac electrical dysfunction and fibrosis, and improvement in survival without overt effects on lung function. Overall, the advent of an immunocompetent model for severe influenza-associated cardiac dysfunction revealed activation of AMPK signaling as a host-targeted metabolic intervention for mitigating virus-induced heart pathologies

## INTRODUCTION

Seasonal influenza viruses cause approximately 1 billion infections annually, resulting in up to 5 million cases of severe illness and 650,000 deaths worldwide^1^. While influenza viruses primarily infect the respiratory tract, cardiac complications, including myocarditis and arrhythmias, occur in up to 40% of hospitalized patients^2, 3, 4, 5, 6^, and 12% experience a major cardiac event such as heart failure^7^. Signs of myocarditis at autopsy have been reported in up to 48% of fatal seasonal influenza cases^4, 8, 9, 10^ and this association is further supported by large-scale epidemiologic studies showing increased cardiovascular events during influenza season, particularly among the unvaccinated^11, 12^. While risk factors for cardiac complications of influenza include advanced age, tobacco use, and diabetes, clinically significant heart damage can also occur in otherwise healthy adults and children^7^. H1N1 influenza A virus infections are most often implicated in these cardiac pathologies, followed by H3N2 and influenza B viruses^13^. As a striking example, an autopsy study of more than 100 individuals during the 1918 H1N1 influenza pandemic revealed severe cardiac injury in nearly all of these fatal cases^14^. Despite the frequency and severity of these complications, no targeted therapeutics exist, and underlying mechanisms remain poorly defined, due in part to a lack of tractable animal models^15,16^.

The three most studied laboratory strains of influenza A virus fail to induce major cardiac phenotypes in immunocompetent mice^17^. Even the most virulent of these, PR8 (H1N1), induces only mild, transient cardiac phenotypes in WT C57BL/6 mice^17, 18, 19^, and the commonly studied WSN (H1N1) and X31 (H3N2) strains produce no significant cardiac pathology^17^. We previously reported that knockout of a single antiviral restriction factor, interferon-induced transmembrane protein 3 (IFITM3), allows increased replication of PR8 in mouse hearts, causing severe arrhythmias, inflammation, and fibrotic remodeling^17^. This system demonstrated that influenza virus can replicate in cardiomyocytes and that viral replication in these cells is required for cardiac pathology^18^. These results challenged the prevailing but weakly supported assumption that cardiac complications of influenza are driven solely by lung-derived systemic inflammation^20^. In contrast, the paradigm that direct viral replication in the heart drives disease is supported by long-standing evidence that influenza virus is cardiotropic, having been detected in human, non-human primate, and rodent hearts^19, 21, 22, 23, 24, 25, 26, 27, 28, 29^. For example, 1918 pandemic H1N1 virus disseminates to the heart by day 3 post infection in Cynomolgus macaques^22^, underscoring the value of animal models for precise temporal analysis of viral dissemination and tissue injury.

Two prior studies of 2009 pandemic H1N1 influenza virus in mice reported cardiac dissemination. In one, obese mice showed increased heart viral loads and modest left ventricular cardiac wall thickening^30^. In another, WT mice infected with a high (10^6^ PFU) dose developed cardiac infection, but rapid lethality limited functional cardiac assessment^31^. These findings align with clinical reports of cardiac complications upon the emergence in of 2009 pandemic H1N1 virus infections in humans and the continued cardiac issues induced by the seasonal descendants of this virus^4, 7, 32, 33^. Recently, our group developed a minimally mouse-adapted 2009 H1N1 variant possessing a single amino acid substitution in the PB2 protein at amino acid position 158 (E158A)^34^. This mutation enabled enhanced lung replication specifically in mice without enhancing replication in human cells^34^. Mutations at this position are thought to modulate viral polymerase activity^35^. We hypothesized that the H1N1-E158A virus may also exhibit enhanced replication in mouse heart tissue, possibly providing an immunocompetent animal model of influenza-associated cardiac disease at physiologically relevant viral doses.

Here we show that the minimally adapted 2009 H1N1 influenza virus disseminates and replicates robustly and persistently in WT mouse hearts, inducing inflammation, electrical dysfunction, structural remodeling, and fibrotic collagen accumulation. Global transcriptomic profiling of influenza virus-infected cardiac tissue revealed distinct early and late gene programs that associate with disease progression. Based on these findings, we demonstrate that pharmacologic activation of AMP-activated protein kinase (AMPK), a key cellular energy sensor^36^, significantly improves cardiac electrical function during infection as well as overall survival. Our findings establish a tractable immunocompetent mouse model for influenza-associated cardiac pathogenesis and identify host-targeted metabolic modulation as a therapeutic approach.

## RESULTS

### Mouse-adapted 2009 H1N1 influenza virus provides a model of severe infection with sustained lung and heart replication

In a previous study, we showed that infection of WT mice by the H1N1-E158A virus at a dose of 10^3^ TCID50 resulted in extreme weight loss and death by day 6 post-infection^34^. The early death at these doses was expected to preclude the study of cardiac complications since heart pathologies do not emerge until day 10 in the *Ifitm3*^-/-^ model of severe influenza^17, 18^. We therefore characterized the effects of a lower dose, 200 TCID50, of H1N1-E158A to determine whether survival times were extended. The lower viral dose continued to provide a severe infection model that induced significantly more weight loss than an equal dose of the parent 2009 H1N1 virus (**Fig 1A**). Importantly, about half of the animals survived to day 10 (**Fig. 1B**), allowing evaluation of lung and heart samples in severe, late-stage disease.

**Figure 1:**
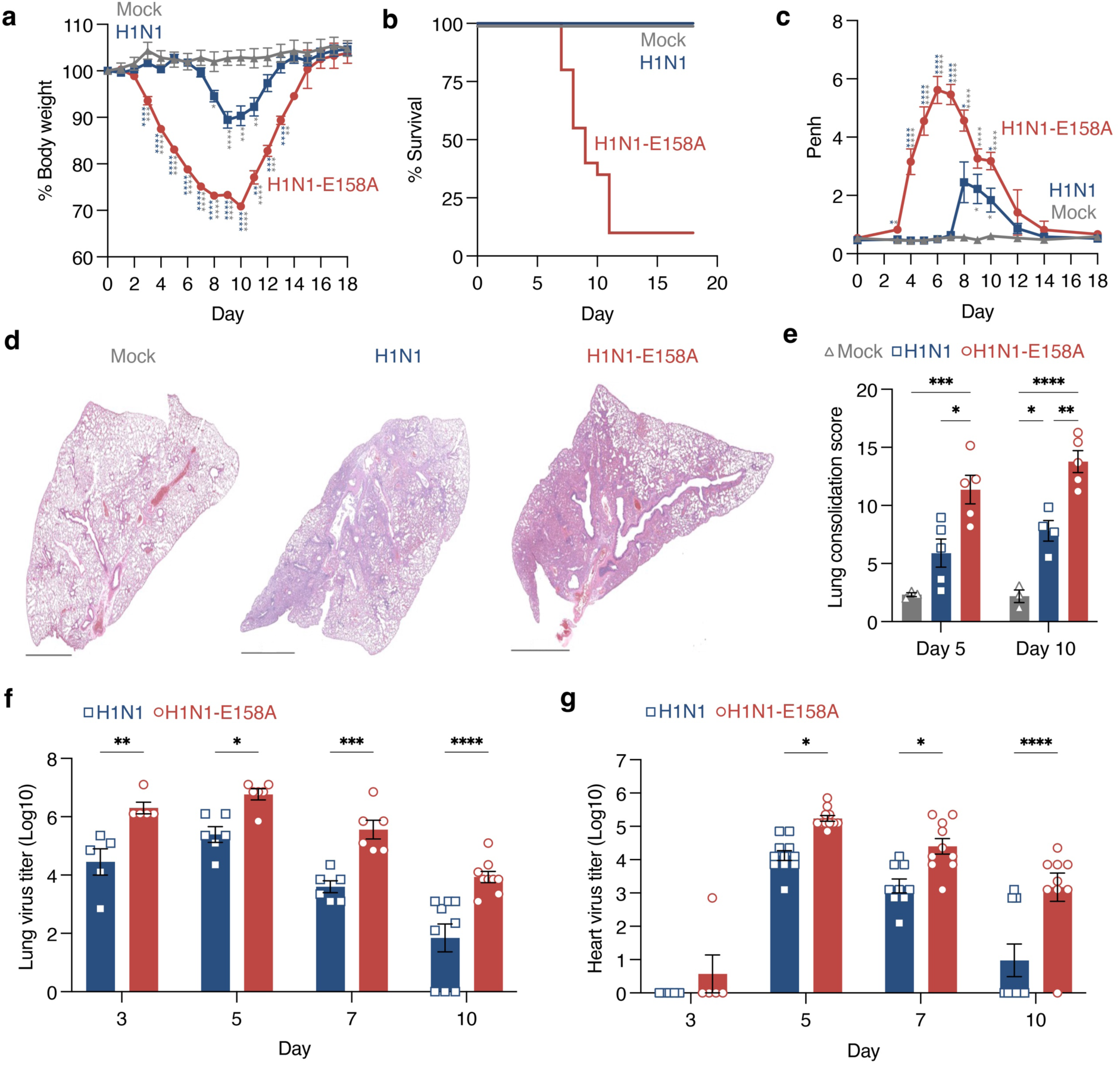
Mouse adapted 2009 H1N1 influenza virus provides a severe infection model with sustained cardiac replication in WT mice. Mice were intranasally inoculated with PBS (mock) or 200 TCID50 of either H1N1 virus or H1N1-E158A virus. **a,** Body weight loss over 18 days. Each data point represents the average weight per mouse normalized to 100% of baseline; error bars indicate SEM. Group sizes: Mock, n = 9; H1N1, n = 16; H1N1-E158A, n = 33. Data are pooled from three independent experiments for mock and H1N1 and four experiments for H1N1-E158A. Statistical significance determined by two-way ANOVA followed by Tukey’s multiple comparisons test; P < 0.05, **P < 0.01, ***P < 0.001, ****P < 0.0001. **b,** Kaplan-Meier survival curves on mice as in **a**. **c,** Whole-body plethysmography measurements of enhanced pause (Penh) as an indicator of respiratory dysfunction. Each point reflects the average of multiple mice; error bars represent SEM. Group sizes: Mock, n = 6; H1N1, n = 12; H1N1-E158A, n = 23. Statistical significance determined by two-way ANOVA followed by Tukey’s multiple comparisons test; P < 0.05, ****P < 0.0001. **d,** Representative H&E-stained lung sections at day 10. Scale bars, 2 mm. **e,** Quantification of lung consolidation (solid tissue vs. open airspace) from whole lung sections at days 5 and 10. Each dot represents lungs from one mouse; error bars indicate SEM. Day 5 p.i.: Mock, n = 3; H1N1, n = 5; H1N1-E158A, n = 5. Day 10 p.i.: Mock, n = 3; H1N1, n = 4; H1N1-E158A, n = 5. Statistical significance determined by two-way ANOVA with Sidak’s multiple comparisons test; *P < 0.05, **P < 0.01, ***P < 0.001, ****P < 0.0001. **f, g,** Viral titers in lung (**f**) and heart (**g**) homogenates measured by TCID50 assay. Each data point represents an individual mouse; error bars indicate SEM. Day 3 p.i.: H1N1 and H1N1-E158A, n = 5 (**f, g**). Day 5 p.i.: n = 6 (H1N1 and H1N1-E158A, f); n = 11 (**g**). Day 7 p.i.: n = 6 (H1N1 and H1N1-E158A, **f**); H1N1, n = 9; H1N1-E158A, n = 10 (**g**). Day 10 p.i.: H1N1 and H1N1-E158A, n = 9 (**f, g**). Statistical significance determined by two-way ANOVA with Sidak’s multiple comparisons test; *P < 0.05, **P < 0.01, ***P < 0.001, ****P < 0.0001.

Lung function measurements via whole body plethysmography indicated earlier and more severe airway obstruction caused by the H1N1-E158A virus in comparison to the parent strain (**Fig 1C**). Accordingly, lung H&E staining showed increased inflammation and tissue consolidation induced by the adapted variant virus at days 5 and 10 post infection (**Fig 1D,E and Supp Fig 1**). These pathologies corresponded to increased lung viral titers for the adapted virus at all timepoints examined (days 3, 5, 7, and 10 post infection) (**Fig 1F**). Moreover, the adapted virus showed significantly higher average heart titers than the parent virus throughout the same time course (**Fig 1G**). Indeed, our data indicate that the parent virus is cleared from the hearts of most mice by day 10 while replication of the E158A adapted variant largely persists through this timepoint (**Fig 1G**). Overall, our results confirm enhanced replication and pathogenesis of the H1N1-E158A variant in the lungs at a dose of 200 TCID50 and newly establish that the variant virus disseminates to the mouse heart, where it establishes a sustained replicative niche.

### H1N1-E158A influenza virus induces significant cardiac electrical dysfunction and structural alterations in WT mice

To assess whether replication of the H1N1-E158A virus in the heart is linked to cardiac electrical phenotypes, we performed electrocardiography (ECG) throughout infection. Both parent 2009 H1N1 virus and the H1N1-E158A variant induced heart rate depression (**Fig 2A**) with corresponding increases in RR (interbeat) intervals (**Fig 2B**) and arrhythmic activity, measured by widened RR interval ranges (**Fig 2C**) and greater RR interval variability (**Fig 2D**) by day 7 post infection. By day 10, mice infected with the parent virus recovered these parameters (**Fig 2C,D**), coinciding with virus clearance (**Fig 1F,G**). In contrast, H1N1-E158A-infected mice maintained their heart rate depression and exhibited strikingly elevated RR interval ranges and variability (**Fig 2C-E**), consistent with marked conduction system dysfunction driven by persistent viral replication in the heart (**Fig 1G**).

**Figure 2:**
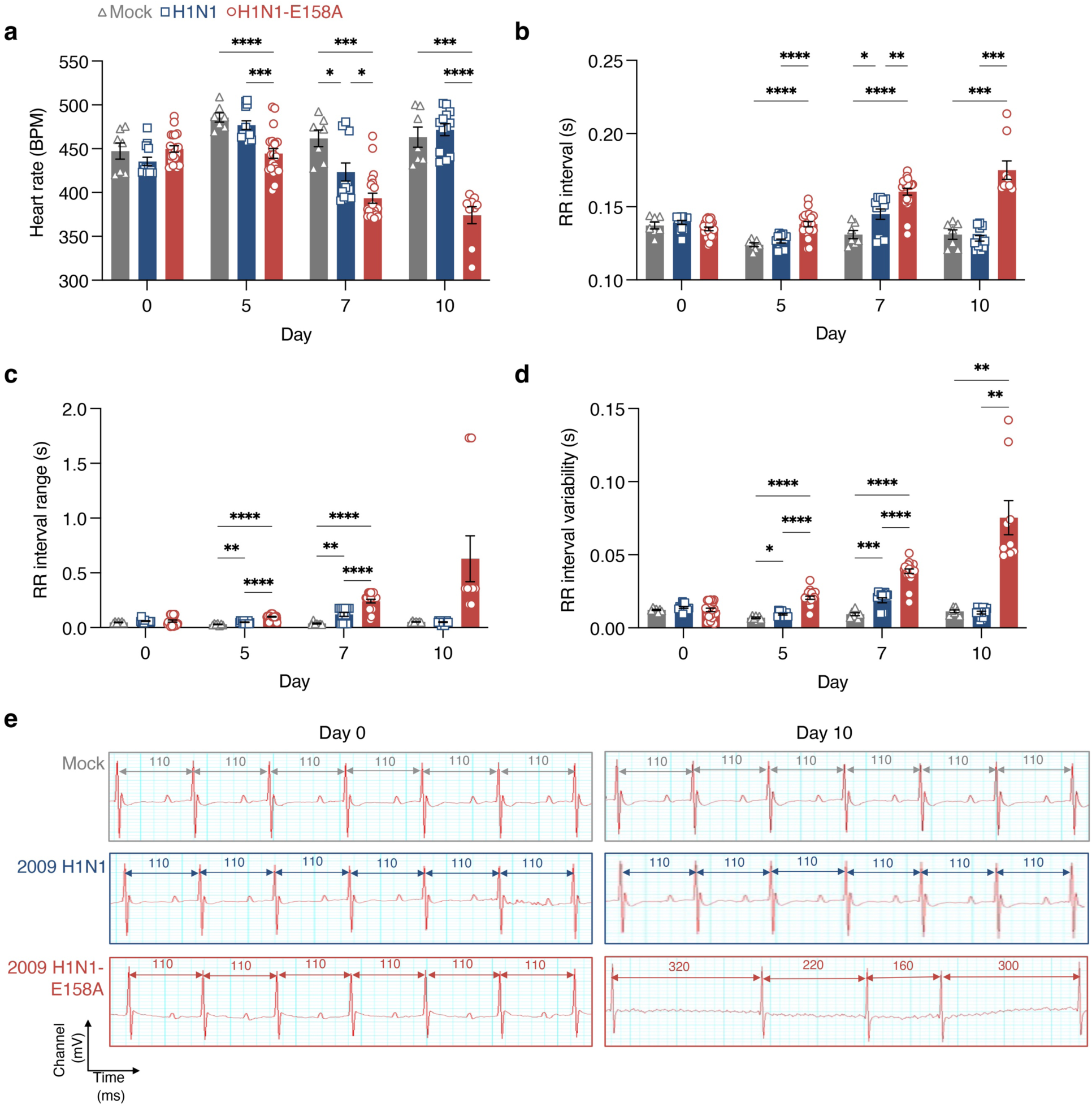
Mouse adapted 2009 H1N1 virus induces cardiac electrical dysfunction in WT mice as measured by electrocardiography. Mice were intranasally inoculated with PBS (Mock) or infected with 200 TCID50 of either H1N1 virus or H1N1-E158A virus. ECG recordings were obtained at defined time points over the course of infection. **a**, Heart rate (beats per minute). **b**, RR interval (interbeat intervals defined as time between R peaks). **c**, RR interval range (defined as the difference between the longest and shortest RR interval during a 5-minute ECG recording). **d**, RR interval variability (defined as the statistical variation in time intervals between among RR intervals in 5-minute ECG recordings. For **a-d**, each data point represents an individual mouse; error bars indicate SEM. Data are pooled from three independent experiments for virus infections and two experiments for Mock (Mock, n = 7; H1N1, n = 13; H1N1-E158A, n = 21, except for Day 10 where n = 9 surviving H1N1-E158A infected animals). Statistical analysis by two-way ANOVA with Tukey’s multiple comparisons test; *P < 0.05, **P < 0.01, ***P < 0.001, ****P < 0.0001. **e**, Representative ECG traces from Mock and virus-infected mice at baseline and 10 days post-infection.

To determine whether the observed electrical abnormalities are accompanied by cardiac structural changes and loss of function, we performed echocardiography during infections with the parent H1N1 virus or H1N1-E158A. The H1N1-E158A virus induced decreased ventricular diameter and volume, as well as decreased stroke volume and cardiac output, phenotypes that were largely absent in infections with the parent virus (**Fig 3A,B**). Ejection fraction and stroke volume were paradoxically increased in the H1N1-E158A-infected hearts, suggesting a compensatory response (**Fig 3B**). Together, these findings demonstrate that the H1N1-E158A virus drives pathological remodeling of the heart, linking enhanced and prolonged cardiac viral replication to structural and functional deterioration, and further establishing this viral strain as a robust model for influenza virus-induced cardiac complications in WT mice.

**Figure 3:**
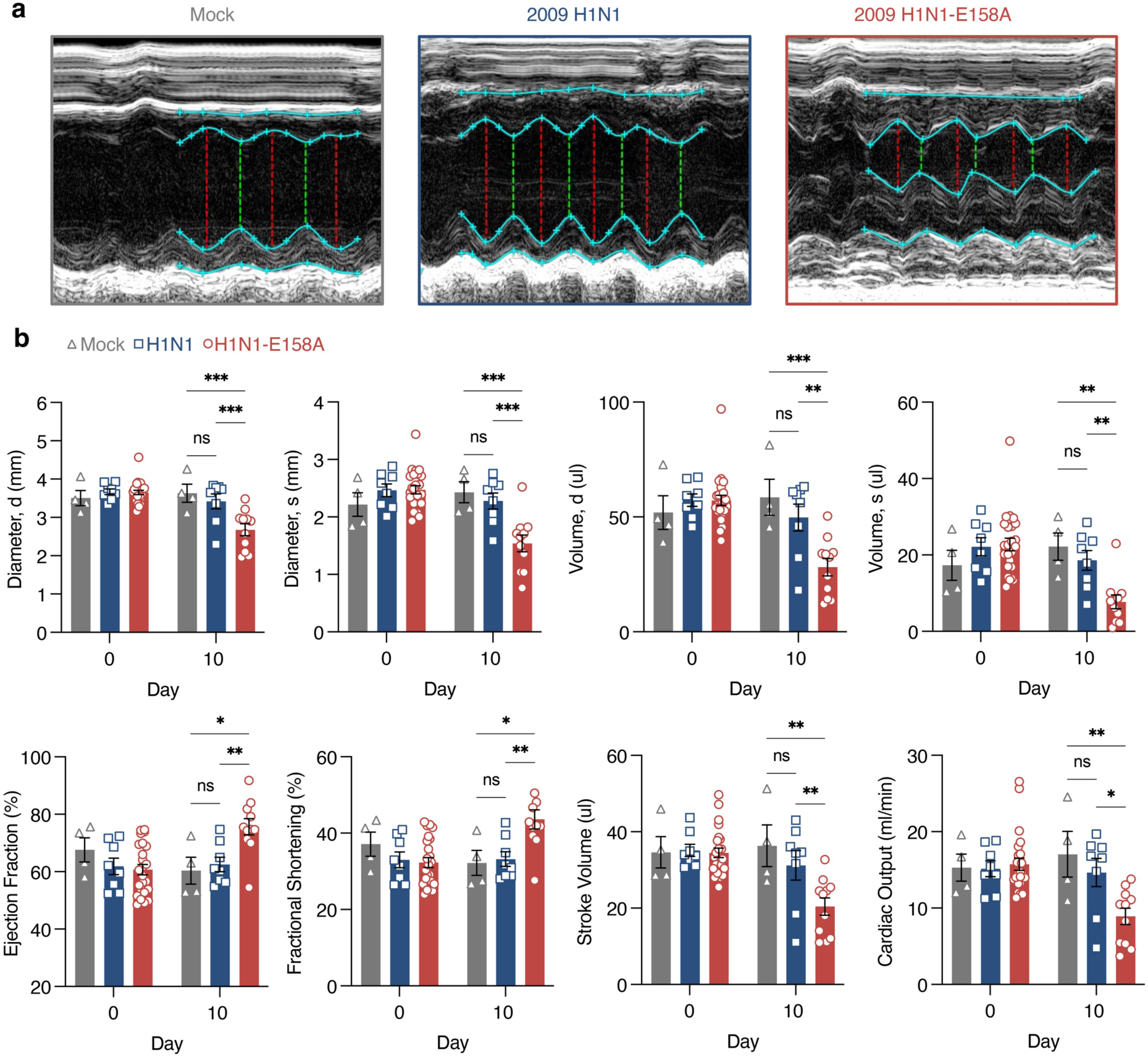
Mouse adapted 2009 H1N1 virus induces cardiac function defects in WT mice as measured by echocardiography. Cardiac morphology and function were evaluated by 2-D echocardiography at baseline or 10 days post mock inoculation with PBS or infection with 200 TCID50 H1N1 virus or H1N1-E158A virus in WT mice. **a**, Representative cardiac images from each group showing left ventricular structure. For each mouse, a minimum of 3 M-mode recordings were analyzed to quantify left ventricular posterior and anterior wall thickness, diastolic and systolic volumes, ejection fraction, fractional shortening, stroke volume, and cardiac output (**b**). Standard echocardiography readout data are plotted with each dot corresponding to an individual mouse. Data represent two independent experiments. Error bars indicate SEM (Pre-infection: Mock n = 4, H1N1 n = 8, H1N1-E158A n = 24; Day 10: Mock n = 4, H1N1 n = 8, H1N1-E158A n = 11 based on animal survival to this timepoint). Statistical significance was determined by two-way ANOVA with Tukey’s multiple comparisons test; *P < 0.05, **P < 0.01, ***P < 0.001.

Given the lethality of the 200 TCID50 dose (**Fig 1B**), we examined whether a lower viral dose would produce similar effects on the heart. A lower dose of the H1N1-E158A virus (100 TCID50) induced less pathogenicity than a dose of 200 TCID50 in terms of weight loss, survival, and lung dysfunction (**Supp Fig 2A-C**). The 100 TCID50 dose also caused detectable heart rate depression and increased RR interval ranges and RR interval variability as measured by ECG (**Supp Fig 2D**). However, the 200 TCID50 dose comparatively induced significantly more severe cardiac electrical dysfunction (**Supp Fig 2D**). Our findings align with observations in humans, where severe cardiac pathologies are most often seen in life-threatening influenza cases requiring hospitalization. Given these results, we continued to use the 200 TCID50 dose as a model for studying severe cardiac disease outcomes but note that lower doses could be used to study mild cardiac manifestations of influenza.

### Heart transcriptomics reveals gene expression changes associated with severe influenza virus infection

To explore the mechanisms of cardiac dysfunction induced by influenza virus, we employed global transcriptomic analysis of H1N1-E158A-infected cardiac tissue in comparison to hearts infected with parent H1N1 virus, providing a range of mild to severe cardiac phenotypes to correlate with gene changes. Specifically, we performed RNA sequencing on heart tissue samples at days 5 and 10 post infection with mock infected animals serving as controls. These timepoints represent an early stage when viral replication is detectable in the heart before the emergence of ECG phenotypes (day 5), and a late stage when cardiac dysfunction is observed in H1N1-E158A infections but has largely resolved for the parent virus (day 10). Sample-to-sample analysis, showed high correlation across the 3-4 biological replicates within each infection group and distinction from mock-infected controls (**Supp Fig 3A**). Multidimensional scaling (MDS) analysis demonstrated distinct transcriptional profiles across treatment groups, with additional separation by virus strain and time point (**Supp Fig 3B**). These results indicate that influenza virus infection elicits consistent, strain-specific transcriptional programs that evolve over time, with maximal divergence from mock controls observed at day 10 for the H1N1-E158A variant.

Consistent with this transcriptional shift, we found that the H1N1-E158A virus induced significantly more differentially expressed genes (DEGs) at both days 5 and 10 post-infection when compared to the parent virus (**Fig 4A-C**). At day 5, the E158A virus induced differential expression of 1266 genes, in the heart when compared with mock infected animals, while the parent virus dysregulated 119 genes (lfc |1|, p-adj = 0.05). At day 10, these differences continued with the E158A virus altering expression of 2873 genes, compared to 458 genes for the parent virus. Clustering and functional enrichment analysis revealed strong induction of adaptive immune pathways, including T cell activation and proliferation, cytokine signaling, and positive regulation of cell adhesion, alongside changes in extracellular matrix organization and mitochondrial respiration **(Fig 4D**). Although both viruses activated similar pathways, the E158A virus infection generally provoked stronger responses involving greater numbers of genes. For example, immune-related networks were more prominent in H1N1-E158A-infected hearts at day 10, suggesting an amplified or prolonged inflammatory response compared to parent virus infection (**Fig 4D**).

**Figure 4:**
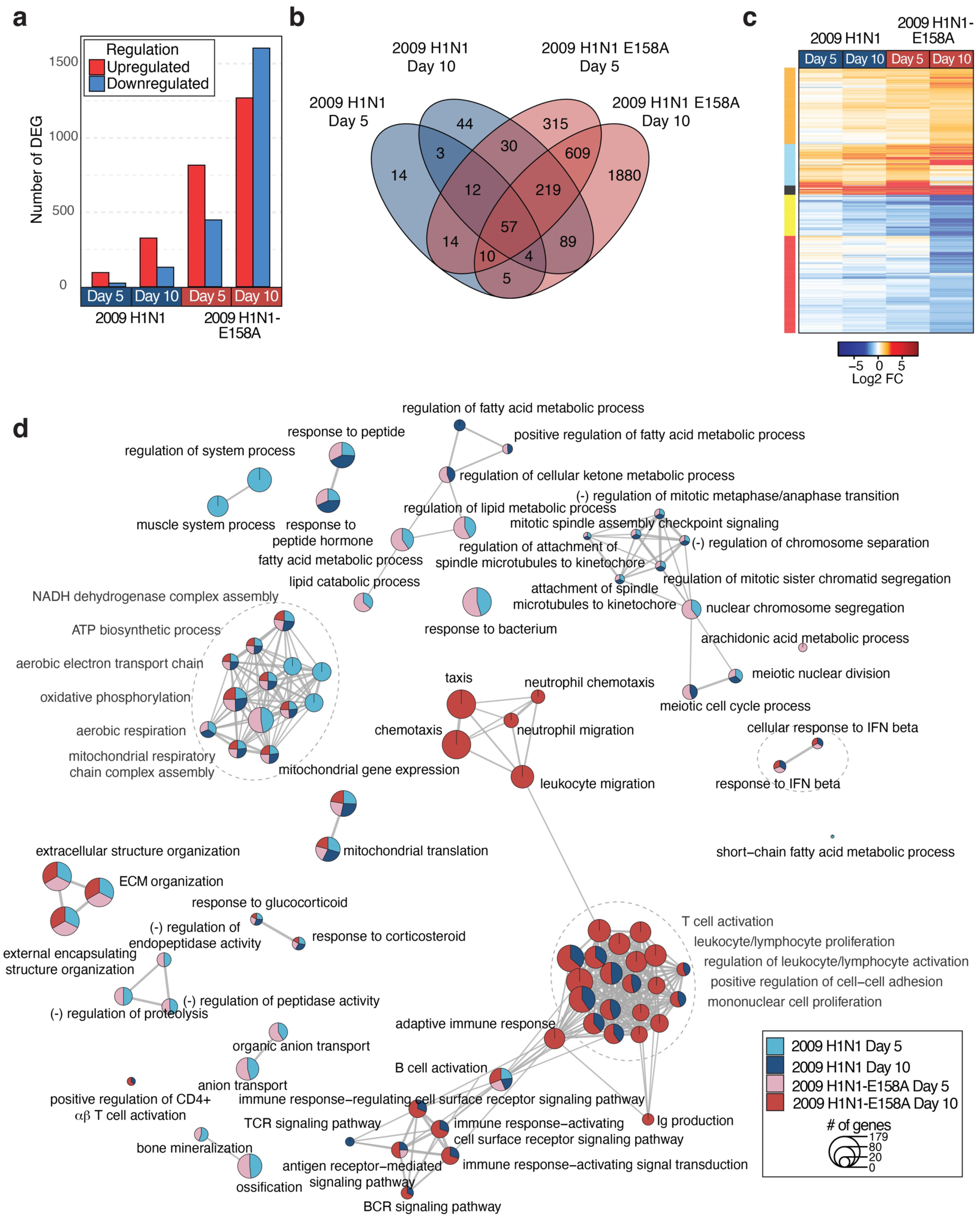
Global transcriptomic changes induced in influenza virus infected hearts. Mice were infected with 200 TCID50 of either H1N1 or H1N1-E158A influenza virus as described above. Total RNA was extracted from cardiac tissue of mock controls (n = 3) and infected mice (n = 4 per group, except day 10 H1N1 n = 3) at days 5 and 10 post-infection and subjected to bulk RNA sequencing to assess host transcriptional responses. **a,** Bar graph of the number of differentially expressed genes (DEGs) identified across various experimental conditions. Color indicates upregulation (red) or downregulation (blue) of genes relative to mock-inoculated mice. Significance cutoff was set at lfc |1| and a p-adjusted value of 0.05. **b,** Venn diagram illustrating the overlap and unique DEGs across different experimental conditions. **c,** Heatmap of the union of DEGs (3305 DEG), highlighting the expression patterns across conditions. DEG were clustered based on Euclidean distances (clusters represented by colors to the left of the heatmap). **d,** Gene Set Enrichment Analysis showing enrichment of Gene Ontology (GO) terms across infectious conditions over time. Node size indicates the number of genes in each GO term. Node color indicates significant enrichment of GO term in the specified condition.

### Mouse-adapted 2009 H1N1 influenza virus induces fibrosis-associated cardiac changes

Gene set enrichment of pathways involved in extracellular matrix remodeling were prominent in H1N1-E158A influenza virus-infected cardiac tissue (**Fig 4D**). Specifically, expression of numerous collagen and matrix remodeling enzymes showed greater alterations by the adapted virus compared to parent (**Fig 5A**), suggesting possible fibrotic remodeling of the heart. We thus examined heart sections from infected and control animals using Masson’s trichrome stain to visualize collagen. We found evidence of fibrotic collagen deposits in H1N1-E158A-infected day 10 hearts with significantly more collagen staining compared to parent virus infected hearts or controls (**Fig 5B,C**). While several collagen genes were downregulated at day 10, the elevated collagen accumulation could possibly be explained by enhanced crosslinking and stabilization of collagen via lysyl oxidase (Lox), which shows a sustained upregulation in H1N1-E158A virus infection (**Fig 5A**). Likewise, Testican-2 (Spock2), a calcium-binding extracellular matrix proteoglycan, was also highly upregulated in H1N1-E158A-infected hearts and may concurrently contribute to increased collagen accumulation through modulation of extracellular matrix architecture and mechanics (**Fig 5A**). Overall, the observed gene expression changes and histological staining indicate that the robust replication of the H1N1-E158A influenza virus causes significant changes in the extracellular matrix of infected hearts.

**Figure 5:**
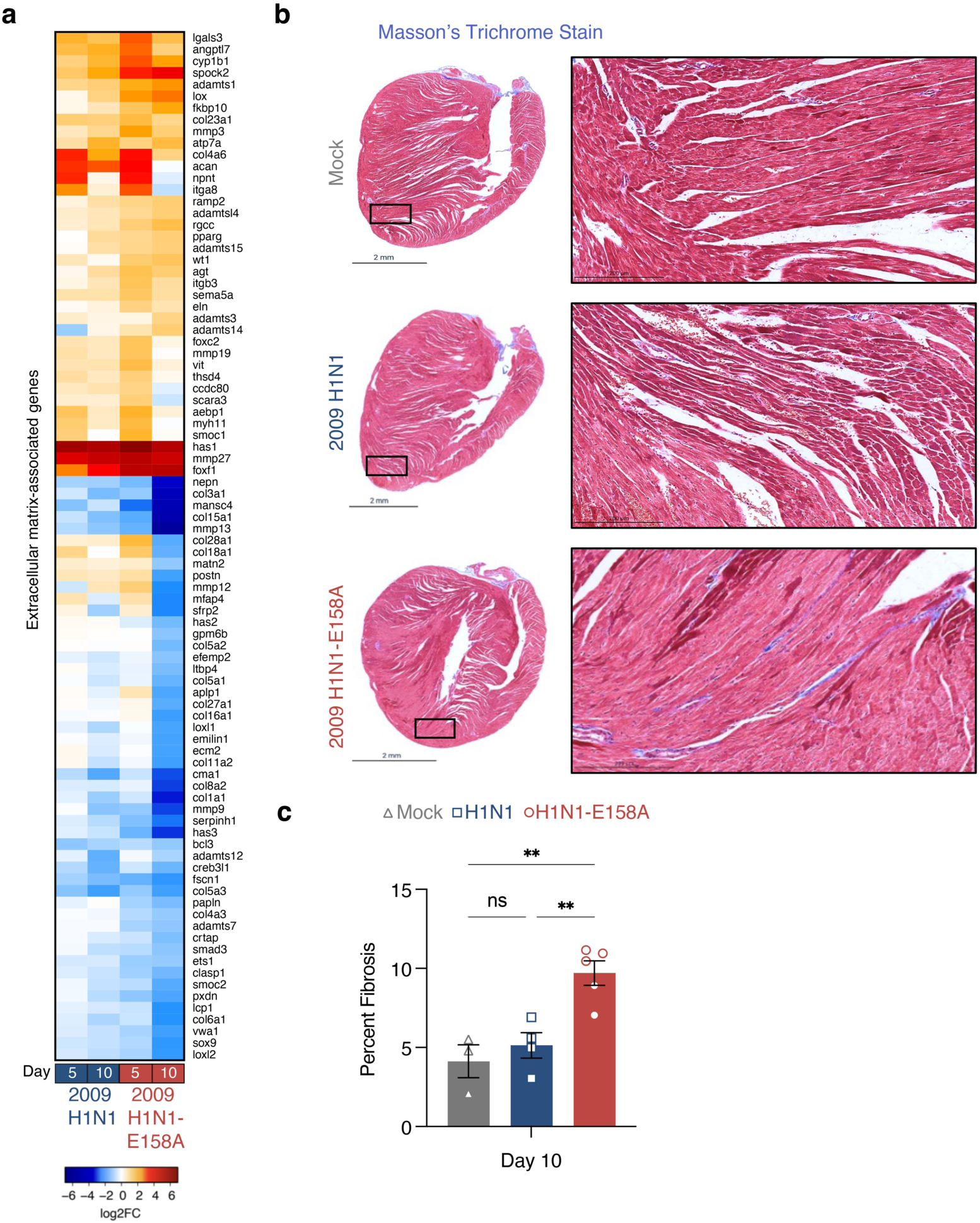
Mouse adapted 2009 H1N1 virus induces cardiac fibrotic remodeling. **a**, Bulk RNA sequencing data from infected mice relative to mock controls (as in Fig. 4) were analyzed to generate a heat map of differentially expressed genes associated with extracellular matrix organization and mechanical properties. **b,** Hearts were harvested 10 days post infection with the indicated virus or mock innoculation and stained with Masson’s trichrome to assess fibrosis (blue staining indicates collagen accumulation). Representative images from each group are shown, with boxed regions magnified at right. **c,** Fibrosis was quantified by calculating the percentage of blue-stained pixels using ImageJ. Each data point represents an individual heart (Mock n = 3, H1N1 n = 4, H1N1-E158A n = 5). Error bars indicate SEM and statistical significance was determined by One-Way ANOVA followed by Tukey’s Multiple Comparisons Test, **P < 0.01.

### Influenza virus induces an inflammatory response in the heart

RNA sequencing results suggested inflammatory and interferon responses are activated by influenza virus infection of the heart (**Fig 4D**). Analysis of 87 interferon stimulated genes (ISGs) identified among the union of DEGs from all infected samples, showed a particularly robust activation of ISG expression by the H1N1-E158A virus with a small subset of genes showing downregulation, particularly in late infection (**Supp Fig 4A**). To further analyze inflammatory responses produced in heart tissue upon infection with H1N1 parent virus or H1N1-E158A, we measured cytokine levels in heart homogenates via ELISA. We found that both parent H1N1 and H1N1-E158A viruses induced cardiac IFNβ, TNF, IL-6, IL-18, and IL-1β above levels seen in mock control animals and that the H1N1-E158A virus induced higher levels of each inflammatory cytokine than the parent virus at day 5 post infection (**Supp Fig 4B**). At days 7 and 10 post infection, cytokine levels were similar in E158A and parent virus-infected samples (**Supp Fig 4B**). The residual inflammation in hearts of parent virus infected animals at day 10 suggests that these particular cytokines are likely not the primary driver of cardiac dysfunction, as these animals have largely resolved their cardiac phenotypes by this timepoint (**Fig 2** and **Fig 3**). Nonetheless, our findings reveal a more robust early immune activation induced by the H1N1-E158A virus in heart tissue.

### AMPK-activating drug AICAR prevents severe influenza-associated cardiac dysfunction

In addition to fibrotic and inflammatory gene changes, we found that a set of mitochondrial-associated genes were downregulated (**Fig 6A**), reinforcing pathway level attenuation (**Fig 4D**). To gain further insights into these transcriptional changes we performed Ingenuity Pathway Analysis, which identified infection-driven changes in canonical pathways related to inflammation, wound healing, circadian rhythm, extracellular matrix, and energy metabolism, among others (**Fig 6B**). Among these, AMPK signaling was prominently activated (**Fig 6B**), indicating that infection induces major metabolic shifts in the heart. Transcriptional regulator analysis also predicted activation of the AMPK downstream effector PGC-1α (PPARGC1A) (**Fig 6C**), a central regulator of mitochondrial biogenesis and fatty acid metabolism^37, 38^. Though perturbation of the AMPK/PGC-1α axis was seen with both parent and H1N1-E158A virus infections, activation was most pronounced on day 10 post infection with the H1N1-E158A virus (**Fig 6B,C**). Supporting a role for early metabolic disruption in heart dysfunction, genes associated with cardiac lipid oxidation were also altered by day 5 post infection (**Supp Fig 5**). Given that cardiomyocytes are the most mitochondria-rich cells in the body and depend heavily on oxidative metabolism for sustained contractility, we reasoned that activation of mitochondrial biogenesis and lipid metabolism via the AMPK/PGC-1α axis may be a compensatory response to infection that could be harnessed for therapeutic benefits if activated earlier in the course of disease.

**Figure 6:**
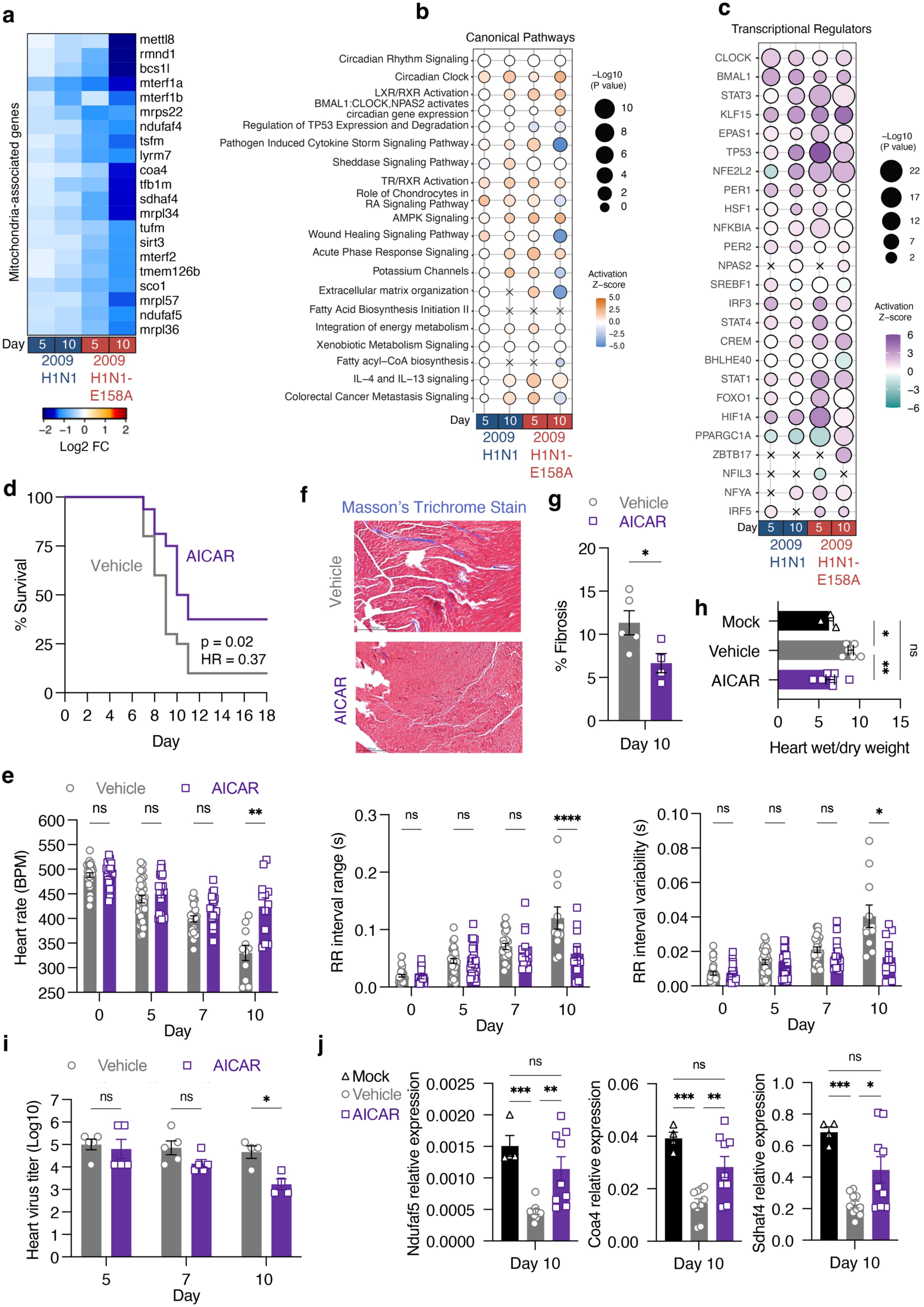
AICAR treatment improves survival and cardiac function during severe influenza virus infection. **a-c,** Bulk RNA sequencing data from infected mice relative to mock controls (as in Fig. 4) were further analyzed. **a**, Heat map of differentially expressed mitochondria-associated genes showing strong infection-induced downregulation particularly by the H1N1-E158A virus at day 10 post infection. **b,** Bubble plot displaying the top 20 enriched canonical pathways identified using Ingenuity Pathway Analysis (IPA). The node size reflects the significance of each pathway (represented by -log10 p-value), while the color indicates the z-score of predicted pathway activation (orange) or inhibition (blue). **c,** Prediction of transcription factor activity based on gene expression profiles using IPA upstream regulator function. Bubble size corresponds to the -log10 p-value of statistical significance, and the color indicates the z-score of predicted activation (purple) or inhibition (teal) of TF activity. **d-j**, To assess therapeutic potential, mice infected with H1N1-E158A (200 TCID50) were treated daily with AICAR (100 mg/kg in saline) or vehicle (saline) from day 4 through day 10 post-infection. **d,** Kaplan-Meier survival curves from two independent experiments (Vehicle: n = 20; AICAR-treated: n = 16). **e,** Representative cardiac sections stained with Masson’s trichrome to visualize collagen (blue) in vehicle- and AICAR-treated mice at day 10 post-infection. **f,** Quantification of cardiac fibrosis from Masson’s trichrome-stained heart sections; each point represents an individual mouse (n = 5 per group). Error bars represent SEM. Statistical comparison by Mann-Whitney test; *\**P = 0.0317. **g,** ECG analysis of heart rate, RR interval range, and RR interval variability in vehicle and AICAR-treated mice over the course of infection. Data are pooled from three independent experiments. Error bars represent SEM. Group sizes: Pre-infection and day 5 post-infection: Vehicle n = 30, AICAR n = 24; Day 7: Vehicle n = 18, AICAR n = 17; Day 10: Vehicle n = 11, AICAR n = 12. Statistical significance determined by two-way ANOVA with Sidak’s multiple comparisons test; *P < 0.05, **P < 0.01, ****P < 0.0001. **h,** Wet/dry weight ratios of hearts collected on day 10 post-infection. Each point represents an individual heart. Error bars represent SEM. Group sizes: Vehicle n =3, AICAR n = 7, Mock n =6. Statistical significance was determined one-way ANOVA followed by Tukey’s multiple comparisons test; *P < 0.05, **P < 0.01. **i,** Viral titers in heart tissue at days 5 and 7 (n = 5 per group), and day 10 (n = 4 per group), determined from a single experiment. Error bars represent SEM. Statistical significance determined by by two-way ANOVA with Sidak’s multiple comparisons test; P = 0.0126. **j,** Relative transcript levels of selected mitochondrial genes in heart tissue from non-infected (Mock), H1N1-E158A infected, and AICAR-treated or vehicle-treated mice at day 10 post infection with 200 TCID50, measured by qRT-PCR. Each data point represents an individual mouse; error bars indicate SEM. Group sizes: Mock, n = 4; H1N1-E158A + AICAR, n = 9; H1N1-E158A + vehicle, n = 10. Data are pooled from two independent experiments. Statistical significance was determined one-way ANOVA followed by Tukey’s multiple comparisons test; *P < 0.05, **P < 0.01, ***P < 0.001.

To test this possibility, we used 5-aminoimidazole-4-carboxamide ribonucleoside (AICAR), a cell-permeable adenosine analog that is phosphorylated intracellularly to form AICAR monophosphate, an AMP mimetic that activates AMPK^39, 40^. Indeed, AICAR has been used to investigate metabolic regulation and tissue protection in sterile ischemic heart injuries in several animal models^36, 41, 42, 43, 44, 45, 46, 47, 48, 49^, and in human trials^50, 51, 52^, though it has not been well investigated for effects in infectious myocarditis. We thus examined whether AICAR treatment of H1N1-E158A virus-infected mice starting at day 4 post infection would affect cardiac outcomes. We observed that AICAR did not significantly change peak weight loss caused by the infection (**Supp Fig 6A**). Similarly, lung function as measured by plethysmography was not improved by AICAR treatment (**Supp Fig 6B**) and lung viral titers were unaffected throughout a time course of infection (**Supp Fig 6C**). However, AICAR treatment significantly improved survival of infected mice. While only 2 of 20 vehicle-treated animals survived, 6 of 16 AICAR-treated mice survived and recovered (**Fig 6D**). Kaplan-Meier analysis revealed that the difference in survival between groups reached statistical significance (log-rank p = 0.02), with a hazard ratio of 0.37 (95% CI: 0.16–0.88), indicating a 63% reduction in the risk of death in AICAR-treated animals (**Fig 6D**).

Importantly, in addition to the survival benefit, cardiac electrical function was markedly improved. Heart rate, which generally declines throughout the first 10 days of infection, was decreased at day 5 and remained largely stable through day 10 in AICAR-treated animals (**Fig 6E**). Similarly, indicators of arrhythmic activity, including RR interval range and variability were significantly improved at day 10 post infection comparing AICAR-treated to vehicle control animals (**Fig 6E**). AICAR treatment similarly normalized infection-induced alterations in key echocardiography parameters, including ventricular diameter, volume, fractional shortening, and ejection fraction at day 10 post infection (**Supp Fig 7**). Similar trends were observed for stroke volume and cardiac output, although these did not reach statistical significance (**Supp Fig 7**).

We also noted significant decreases in fibrotic collagen staining in hearts from mice treated with the drug (**Fig 6F,G**), consistent with reported roles of AMPK in limiting cardiac fibrosis^49, 53^. In parallel, we found that infection increased fluid accumulation in the heart, as reflected by an elevated wet-to-dry weight ratio, and that AICAR treatment normalized this to levels comparable to mock-infected animals (**Fig 6H**).

In contrast to a lack of effect on viral titers in the lung, AICAR treatment decreased viral titers in the heart at day 10 post infection (**Fig 6I**). Given that AICAR treatment starting on day 4 did not affect cardiac viral titers at days 5 or 7 in the heart, this suggests that AICAR did not affect viral dissemination to this organ but did elicit an environment in the heart that is less permissive to influenza virus replication by day 10 (**Fig 6I**). AICAR treatment largely normalized expression of tested mitochondria-associated genes that were downregulated in infected control animals (*Ndufaf5*, *Coa4*, and *Sdhaf4*, **Fig 6A**) and showed that AICAR treatment largely normalized expression of these genes (**Fig 6J**). We conclude that AICAR lessens cardiac electrical dysfunction, fibrotic responses, mitochondria-associated gene dysregulation, and overall mortality, supporting the AMPK pathway as a viable therapeutic target in severe influenza virus infections involving the heart.

## DISCUSSION

We describe here a novel, immunocompetent mouse model of severe influenza-associated cardiac dysfunction using a minimally adapted 2009 H1N1 variant carrying a single amino acid substitution in the PB2 polymerase subunit (E158A)^34^. This adaptation enhances viral replication in mouse lung and heart tissue, enabling sustained cardiac infection required for measurable functional and structural heart pathology in WT C57BL/6 mice. Importantly, the PB2 E158A substitution, while boosting replication in murine cells and tissues, reduces replication efficiency in human respiratory epithelial cells, indicating a host-specific trade-off that enhances model safety^34^. Unlike most existing influenza mouse models, which require immunodeficiency or produce only mild and transient cardiac effects, this model recapitulates hallmarks of severe clinical disease, including conduction system abnormalities and reduced cardiac output^7^. These features establish the H1N1-E158A model as a tractable and physiologically relevant system for mechanistic dissection of viral cardiac pathogenesis and preclinical therapeutic testing.

Transcriptomic analyses from this model revealed robust but late activation of the AMPK/PGC-1α metabolic axis, a stress-responsive pathway critical for mitochondrial homeostasis and energy balance, and enabled us to demonstrate that pharmacologic AMPK activation confers significant cardioprotection. Thus, this model not only recapitulates key features of severe disease but also provides a powerful platform to identify and evaluate host-targeted therapeutic strategies.

Our findings reinforce the paradigm that direct viral replication in the heart is a key driver of cardiac complications during influenza virus infection. Persistent replication of the H1N1-E158A virus coincided with sustained heart rate depression, extreme arrhythmic electrical activity, and reduced ventricular function, whereas the parent H1N1 virus was largely cleared from the heart by day 10, allowing modest cardiac phenotypes to resolve. These data challenge the notion that influenza-associated cardiac injury is primarily mediated by systemic inflammation derived from the lung^54^. Instead, our results align with clinical and experimental evidence that influenza viruses are cardiotropic pathogens capable of infecting and damaging cardiomyocytes^17, 18, 19, 21, 22, 23, 24, 25, 26, 27, 28, 29, 30, 31, 55^.

Transcriptomic analysis of infected cardiac tissue revealed broad and sustained transcriptional reprogramming in the H1N1-E158A group, including strong induction of immune and interferon-stimulated gene networks, extracellular matrix remodeling pathways, and metabolic signaling programs. Both parent H1N1 and H1N1-E158A viruses activated similar functional pathway categories, but the magnitude and persistence of these responses were greater in the H1N1-E158A group. Notably, mitochondrial metabolism pathways and circadian rhythm regulators were disrupted alongside inflammatory signatures, identifying energetic stress and disruption of intrinsic cardiac timing mechanisms as additional unexpected consequences of heart infection.

Among the most prominent molecular changes was the activation of the AMPK/PGC-1α axis, which is known to promote mitochondrial biogenesis^37, 56^ and preserve cardiac function under stress^57^. However, activation of this axis peaked at day 10 in the E158A group, well after the onset of electrical dysfunction, and did not correspond to recovery of affected mitochondria-associated genes. This temporal pattern is consistent with a delayed or insufficient compensatory response, in which the heart attempts to restore mitochondrial capacity after substantial injury has already occurred, providing us with a rationale to test whether early pharmacologic AMPK activation could mitigate cardiac damage.

Treatment with the AMP analog AICAR, initiated on day 4 post infection, significantly improved survival, preserved cardiac electrical stability, reduced fibrosis, and restored expression of mitochondria-associated genes in E158A-infected mice. Of note, these benefits were observed without improvement in lung function or reduction in pulmonary viral titers, indicating that cardiac protection was not simply a consequence of systemic antiviral effects. Instead, our data point to a tissue-specific impact of AICAR on influenza virus replication. While early cardiac seeding was unaffected (viral titers at days 5 and 7 post infection were not statistically different between treated and control animals), AICAR significantly reduced viral burden in the heart at day 10. These findings suggest that AMPK activation modifies the cardiac microenvironment in ways that restrict viral replication, highlighting the context-dependent consequences of metabolic modulation. Prior work demonstrated that prophylactic AICAR treatment, initiated three days before infection, reduced weight loss and improved survival in influenza virus-infected mice^58^. However, they did not measure viral titers or assess tissue-specific outcomes but hypothesized that AICAR downmodulated detrimental inflammatory responses. In contrast, our study tested AICAR in a therapeutic setting and identified a selective beneficial effect in the heart, underscoring the potential of AMPK activation as a host-directed, organ-targeted strategy for mitigating fatal cardiac complications of severe influenza.

Consistent with these concepts, previous studies have demonstrated antiviral effects of AICAR through AMPK activation against hepatitis C virus^59^, coxsackievirus^60^, and human herpesvirus 6A^58, 61^, that were largely attributed to metabolic alterations. However, AMPK may be beneficial to the virus in other systems, such as infections with hepatitis B virus^62^ or Ebola virus^63^. Together, the published work on AMPK in virus infections along with our results on influenza virus in the lung and heart suggests that AMPK regulates infection outcomes in a virus- and tissue-specific manner^64^.

The cardioprotective effect of AICAR in our model is consistent with its established role in experimental cardiac ischemia-reperfusion injury^36, 41, 42, 43, 44, 45, 46, 47, 48, 49^ and underscores the potential for AMPK activation as a therapeutic strategy in infectious cardiomyopathies. Although AICAR is a valuable experimental tool, its clinical use is limited by low oral bioavailability and mild safety concerns, such as hyperuricimea^65^, and it is prohibited in competitive sports as a performance-enhancing agent. Nevertheless, given the absence of targeted options for influenza-associated cardiac complications, short-term AMPK activation merits further evaluation, particularly where benefits may outweigh manageable side effects. Other clinically available agents such as metformin, which engages AMPK-dependent and AMPK-independent pathways and has a favorable safety profile, also warrant investigation in this context.

In summary, the H1N1-E158A virus model enables robust and reproducible study of influenza virus-induced cardiac dysfunction in immunocompetent mice, recapitulating key clinical features of severe disease. Using this model, we identify dysregulated cardiac metabolism as a component of viral pathogenesis and demonstrate that AMPK activation via AICAR offers significant protection against cardiac injury. Future work will determine whether this protective effect extends across diverse host backgrounds, repeated infection scenarios, and other cardiotropic viral infections. Overall, our findings establish both a new model for the field and a proof-of-principle for host-targeted metabolic modulation as a strategy to prevent fatal cardiac outcomes in influenza and other cardiotropic infections.

## MATERIALS AND METHODS

### Virus Stocks and Virus Titers

The influenza A virus strains used in this study included A/California/04/2009 (H1N1; BEI Resources, NR-13659) and a mouse-adapted variant carrying a glutamic acid to alanine substitution at position 158 in the PB2 segment (H1N1-E158A), as previously described^34^. Both viruses were propagated for 48 hours in mycoplasma-free Madin-Darby Canine Kidney (MDCK) cells (BEI Resources, NR-26-28) in the presence of TPCK-treated trypsin (Worthington Biochemical Corporation, LS02123). Virus stocks were flash-frozen and stored at -80 °C for single-use applications. To assess viral replication *in vivo*, heart and lung tissues were harvested at designated time points, homogenized in 1 ml phosphate-buffered saline (PBS), flash-frozen, and stored at -80 °C until viral titering on MDCK cells.

### Mice, Infection, and Drug Treatment

Female wild-type C57BL/6J mice (8–10 weeks old) were obtained from Jackson Laboratories (strain #000664) and housed in an ABSL-2 facility at The Ohio State University Biomedical Research Tower. The vivarium was maintained at 20-24 °C (68-76 °F), with a 12:12 h light–dark cycle and relative humidity of 30–70%. Mice were housed in autoclaved, individually ventilated cages (Allentown) containing 0.25 inch of corn cob bedding (Bed-oCobs, The Andersons) and cotton square nesting material. Animals had unlimited access to irradiated, natural ingredient chow (Envigo Teklad Diet 7912) and reverse osmosis-purified water delivered via an automated rack system.

At the time of infection, mice were age-matched within each experimental group. These experiments were not blinded and animals were randomly assigned to treatment groups. Mice were anesthetized with 3% isoflurane in oxygen (flow rate: 1.0 mL/min) and intranasally inoculated with 200 TCID50 of wild-type H1N1 or either 100 or 200 TCID50 of the H1N1-E158A variant, delivered in 50 μL sterile clinical grade saline.

For therapeutic intervention studies, animals received daily intraperitoneal injections of AICAR (100 mg/kg; MedChemExpress, HY-13417) diluted in 200 μL of saline, or an equivalent volume of vehicle alone, from day 4 to day 10 post-infection. In all mouse experiments, clinical signs, including body weight, were monitored daily throughout the course of infection. Mice reaching greater than 30% weight loss with loss of ambulation were humanely euthanized. All procedures were conducted in accordance with protocols approved by The Ohio State University Institutional Animal Care and Use Committee (IACUC protocol number 2016A00000051-R3) and adhered to established ethical guidelines for animal research.

### Whole Body Plethysmography

Lung function was evaluated using whole-body plethysmography (Buxco Small Animal Whole Body Plethysmography system, DSI Inc.). Mice were acclimated to the plethysmography chambers for 10 minutes per day over three consecutive days prior to infection. Respiratory measurements were recorded at baseline and daily throughout infection. At each designated time point, mice underwent a 5-minute acclimation period within the chamber, followed by a 5-minute data acquisition phase. Enhanced pause (PenH) and other plethysmographic parameters were recorded at 10-second intervals throughout the 5-minute reading and subsequently averaged to yield daily individual measurements as reported previously^66^.

### Lung and Heart Histology

For histological analysis, mice were assigned to predetermined harvest days prior to the initiation of experiments. The left lung lobe was excised, fixed in 10% neutral-buffered formalin (Azer Scientific, Inc. Morgantown, PA) at 4 °C for 24 hours, then transferred to 70% ethanol before paraffin embedding and sectioning. Lung sections were stained with hematoxylin and eosin (H&E) and imaged by Histowiz (Histowiz.com, Brooklyn, NY, USA). Hearts were perfused with 5 mL ice-cold 1x PBS prior to careful dissection from the thoracic cavity for fixation in 10% neutral-buffered formalin and subsequent Masson’s Trichrome staining performed by Histowiz. Unbiased quantitative analysis of the stained lung and heart sections was performed using ImageJ software employing the color deconvolution method^67, 68, 69^.

### Electrocardiography

Mice were anesthetized as previously described and positioned prone on a heated pad to maintain normothermia throughout the procedure. Anesthesia was sustained for the duration of data acquisition. Three subcutaneous electrodes were placed under the skin in a lead II configuration to record subsurface electrocardiograms (ECGs). ECG signals were continuously recorded for 5 minutes using a PowerLab 4/30 data acquisition system (AD Instruments, Houston, TX). Recorded traces were subsequently analyzed using LabChart 9 Pro software (AD Instruments).

### Echocardiography

Two-dimensional echocardiography was performed on infected and control mice at baseline and 10 days post-infection using the Vevo 2100 system (VisualSonics). Following anesthesia, mice were placed prone on a heated pad, and chest hair was removed with depilatory cream to optimize acoustic coupling. Proper anatomical orientation was achieved by positioning the transducer along the parasternal long axis at the level of the papillary muscles. Images were acquired and analyzed in M-mode to assess cardiac function, including ejection fraction, fractional shortening, ventricular dimensions, and left ventricular wall thickness.

### ELISA Cytokine Measurements

Mouse IFN-β (DY8234-05), IL-1β (DY401), TNF (DY410), IL-6 (DY406), and IL-18 (DY7625-05) cytokine levels in cardiac homogenates were quantified using ELISA kits (R&D Systems) following the manufacturer’s protocols.

### RNA Sequencing and Analysis

Total RNA was extracted from cardiac tissue homogenates of infected and uninfected mice at baseline and 10 days post-infection with TRIzol (Invitrogen) using the phenol-chloroform extraction method. Library preparation and RNA sequencing were performed by GENEWIZ (GENEWIZ LLC/Azenta US, Inc., South Plainfield, NJ, USA) following their standardized protocols as stated: “The RNA samples received were quantified using Qubit 2.0 Fluorometer (ThermoFisher Scientific, Waltham, MA, USA) and RNA integrity was checked using TapeStation (Agilent Technologies, Palo Alto, CA, USA). The RNA sequencing libraries were prepared using the NEBNext Ultra II RNA Library Prep Kit for Illumina using manufacturer’s instructions (New England Biolabs, Ipswich, MA, USA). Briefly, mRNAs were initially enriched with Oligod(T) beads. Enriched mRNAs were fragmented for 15 min at 94°C. First strand and second strand cDNA were subsequently synthesized. cDNA fragments were end repaired and adenylated at 3′ends, and universal adapters were ligated to cDNA fragments, followed by index addition and library enrichment by PCR with limited cycles. The sequencing libraries were validated on the Agilent TapeStation (Agilent Technologies, Palo Alto, CA, USA), and quantified by using Qubit 2.0 Fluorometer (ThermoFisher Scientific, Waltham, MA, USA) as well as by quantitative PCR (KAPA Biosystems, Wilmington, MA, USA). The sequencing libraries were multiplexed and clustered onto a flowcell. After clustering, the flowcell was loaded onto the Illumina HiSeq instrument according to manufacturer’s instructions. The samples were sequenced using a 2 x 150 bp Paired End (PE) configuration. Image analysis and base calling were conducted by the HiSeq Control Software (HCS). Raw sequence data (.bcl files) generated from Illumina HiSeq was converted into fastq files and de-multiplexed using Illumina bcl2fastq 2.17 software. One mis-match was allowed for index sequence identification.” Raw fastq files were processed, aligned, and quantified with using ROSALIND (ROSALIND, Inc. San Diego, CA, https://rosalind.bio/) as described previously^66, 68, 70^. Correlation-based outlier detection was used to identify outliers (Z-score >2) prior to statistical analysis and excluded from downstream processing. One statistical outlier from the H1N1 day 10 group was removed from further analysis (sample #4). Normalization and statistical analysis for differential gene expression were performed using DESeq R library. Gene Ontology and functional enrichment analysis was done with clusterProfiler (v4.14.16)^71^ and Ingenuity Pathway Analysis (Qiagen). Data visualization was performed in R (4.4.2). Raw data files have been uploaded to the GEO database and will be released upon journal publication.

### Quantitative PCR

RNA was extracted from cardiac homogenates of non-infected, AICAR-treated infected, and vehicle treated infected mice as described above. Complementary DNA (cDNA) synthesis was performed using the AffinityScript qPCR cDNA Synthesis Kit (Agilent; catalogue no. 600559) according to the manufacturer’s instructions. Quantitative PCR (qPCR) was conducted using iTaq Universal SYBR Green Supermix (Bio-Rad) on a CFX384 Real-Time PCR Detection System (Bio-Rad). Gene expression levels were normalized to the reference gene *Chmp2a* using the primer sequences: forward -AGA CGC CAG AGG AAC TAC TTC and reverse -ACC AGG TCT TTT GCC ATG ATTC. Other primers used in this study were acquired from Integrated DNA Technologies and include: *Ndufaf5*: forward -CTT GGG ACA TGT GGT CTC TAA C, and reverse - AGT CAG GAC ATC CAG CAA AC, *Coa4*: forward -GAG AGA CCT GGG AAG AAG AAA C, and reverse - TTG GGT CTG GCT GTC AAT C, *Sdhaf4*: forward - GAG AGC CAC TGC AGA AGT TT, reverse -TCC CAG TCT CCA TAT CGT GTA G. All primer sequences are presented in the 5’ to 3’ orientation.

### Statistics

Data visualization and statistical analyses were performed using built-in functions in GraphPad Prism (v10.5.0). Specific statistical tests and the number of animals used in each experiment are detailed in the corresponding figure legends. A *P* value of <0.05 was considered statistically significant.

## ACKNOWLEDGEMENTS

The authors thanks Dr. Eugene Oltz for helpful discussion and critical review of the manuscript. This work was supported in part by NIH grants R01AI130110 and R01HL157215 to J.S.Y., a Burroughs Wellcome Fund Investigators in the Pathogenesis of Infectious Diseases Award to A.F., American Heart Association Predoctoral Fellowships to J.R. and M.M., and a postdoctoral fellowship from the Interdisciplinary Program in Microbe-Host Biology supported by NIH training grant T32AI165391 to J.C.

---

**Supp Figure 1:**
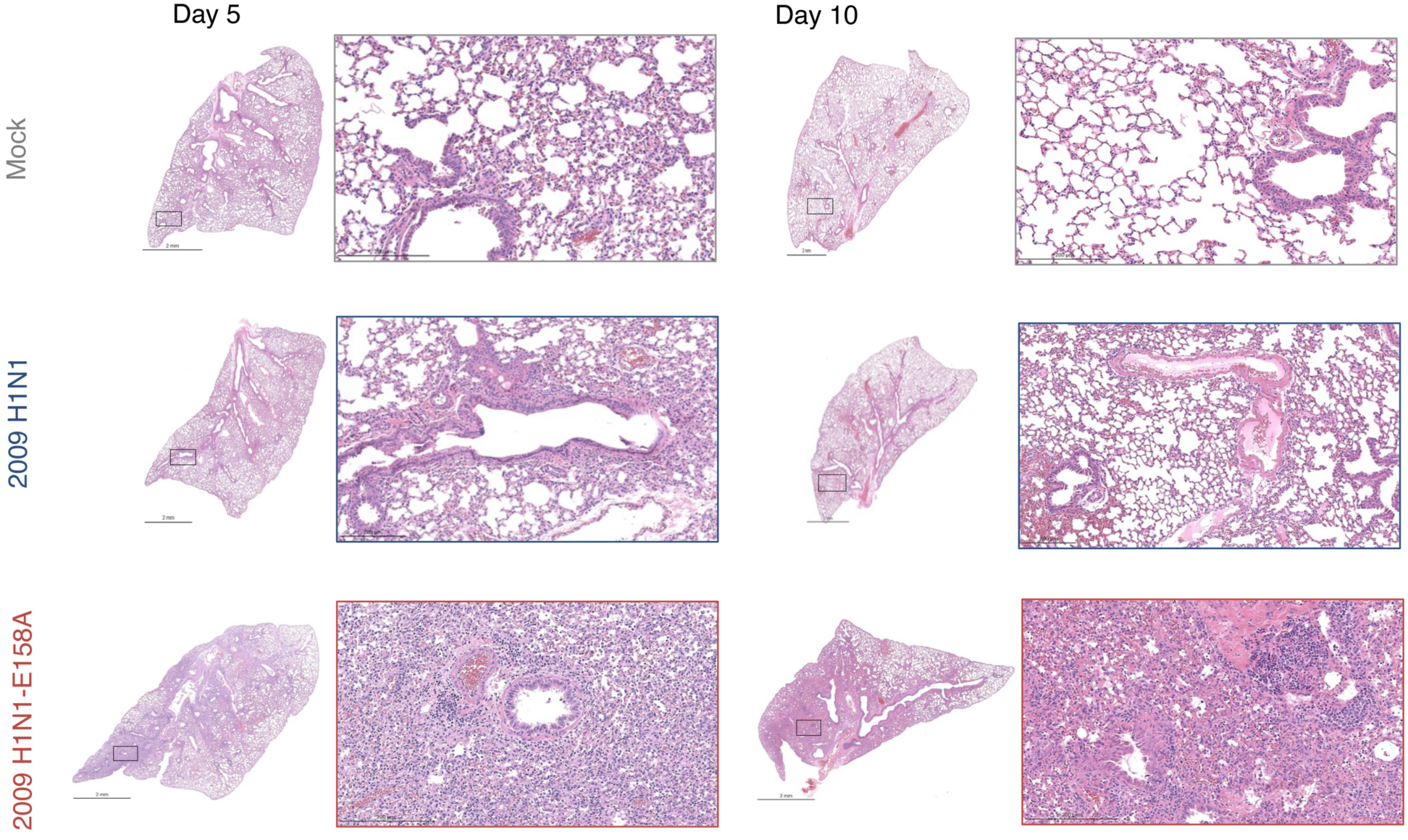
Mouse adapted H1N1-E158A virus robustly induces lung histological damage. Representative H&E-stained lung sections at days 5 and 10 post-infection with 200 TCID50 H1N1 virus or H1N1-E158A virus or mock (PBS) innoculation. Whole-lung sections (left panels; scale bars, 2 mm) and corresponding high-magnification images (right panels; scale bars, 200 μm) are shown for each condition and time point. Boxes in whole-lung images indicate the regions displayed at higher magnification. Note: Day 10 unzoomed images are repeated from main text Figure 1D.

**Supp Figure 2:**
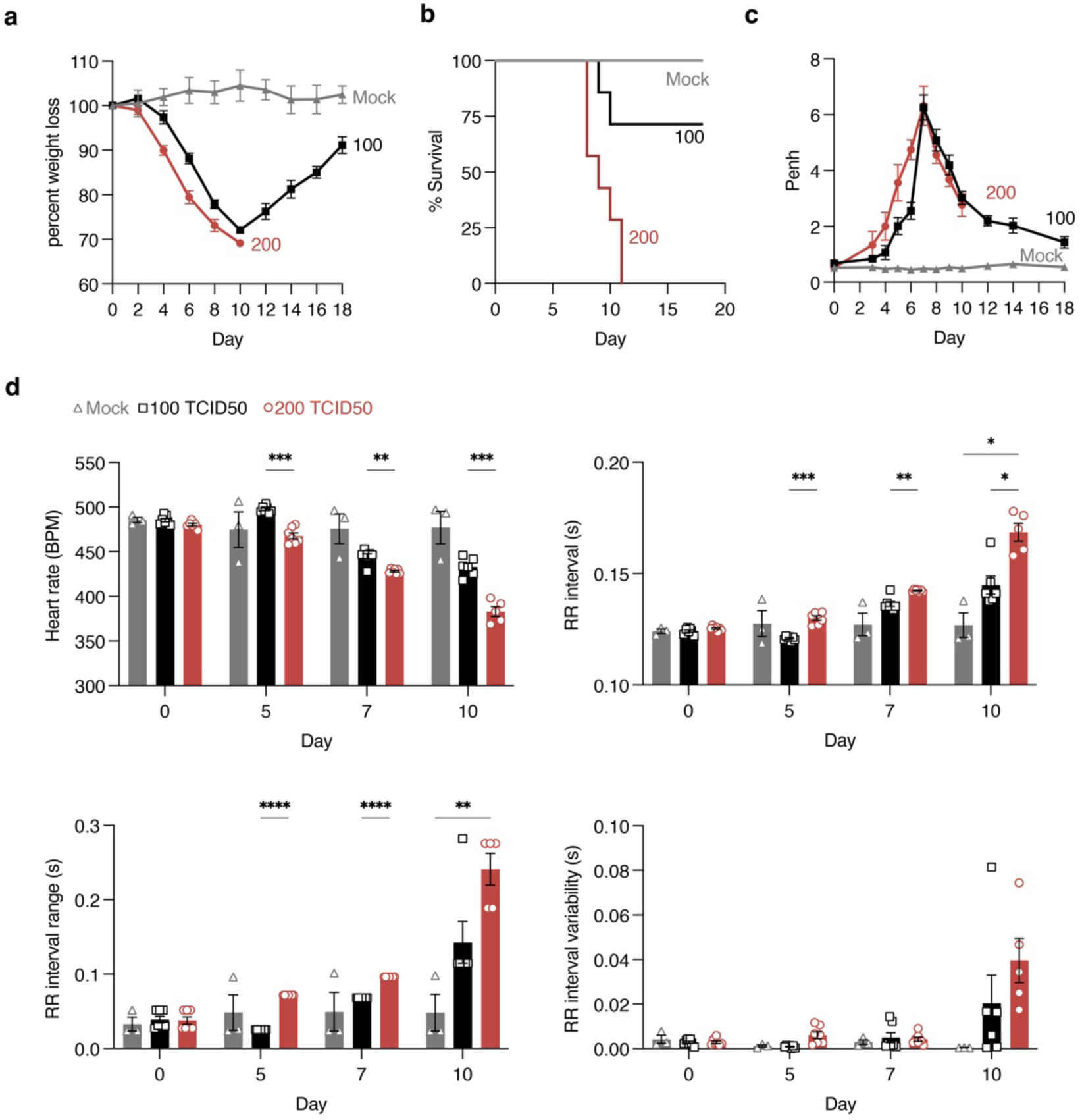
Viral dose responsiveness of lung and heart functional readouts. Mice were intranasally infected with 100 or 200 TCID50 of H1N1-E158A virus or were mock (PBS) innoculated. **a,** Body weight loss following infection. Each data point represents the average weight of individual mice normalized to 100% of their starting weight on day 0. Error bars indicate SEM. **b,** Kaplan-Meier survival curves. **c,** Whole-body plethysmography analysis of Penh as an indicator of respiratory dysfunction. Each data point represents the average Penh value from multiple mice; error bars indicate SEM. Mice in **a,b,c** were monitored daily for 18 days. Group sizes: Mock, n = 3; H1N1, n = 7; H1N1-E158A, n = 7. **d,** Heart rate, RR interval, RR interval range, and RR interval variability in Mock and H1N1-E158A infected mice at baseline and at days 5, 7, and 10 post-infection. Group sizes: Mock, n = 3 (all time points); H1N1-E158A (100 TCID50): n = 7 (baseline, days 5 and 7), n = 6 (day 10); H1N1-E158A (200 TCID50): n = 7 (baseline, days 5 and 7), n = 5 (day 10). All data are from a single experiment. Statistical analysis by two-way ANOVA with Sidak’s multiple comparisons test; *P < 0.05, **P < 0.01, ***P < 0.001, ****P < 0.0001.

**Supp Figure 3:**
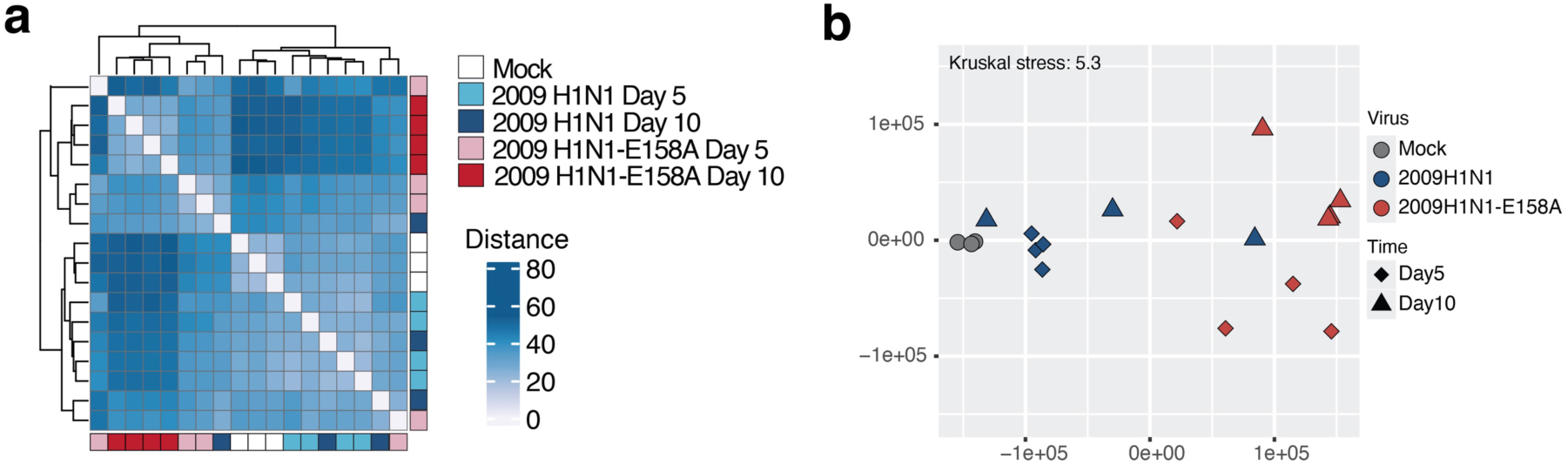
RNA sequencing gene expression correlation and principal components analysis for individual samples. Bulk RNA sequencing was performed on heart samples from mice infected with 200 TCID50 H1N1 virus or H1N1-E158A virus in comparison to mock (PBS) inoculated animals. **a,** Heatmap showing the correlation of gene expression profiles across all samples. The color scale indicates the degree of similarity between samples. **b,** Multidimensional scaling (MDS) representation of the heart transcriptional responses elicited by H1N1 and H1N1-E158A influenza virus strains at days 5 and 10. Each data point represents a biological replicate, where color denotes viral infection and shape indicates timepoint post infection. The quality of the representation is provided by the Kruskal stress value, with a low percentage of Kruskal stress (5.3%).

**Supp Fig 4:**
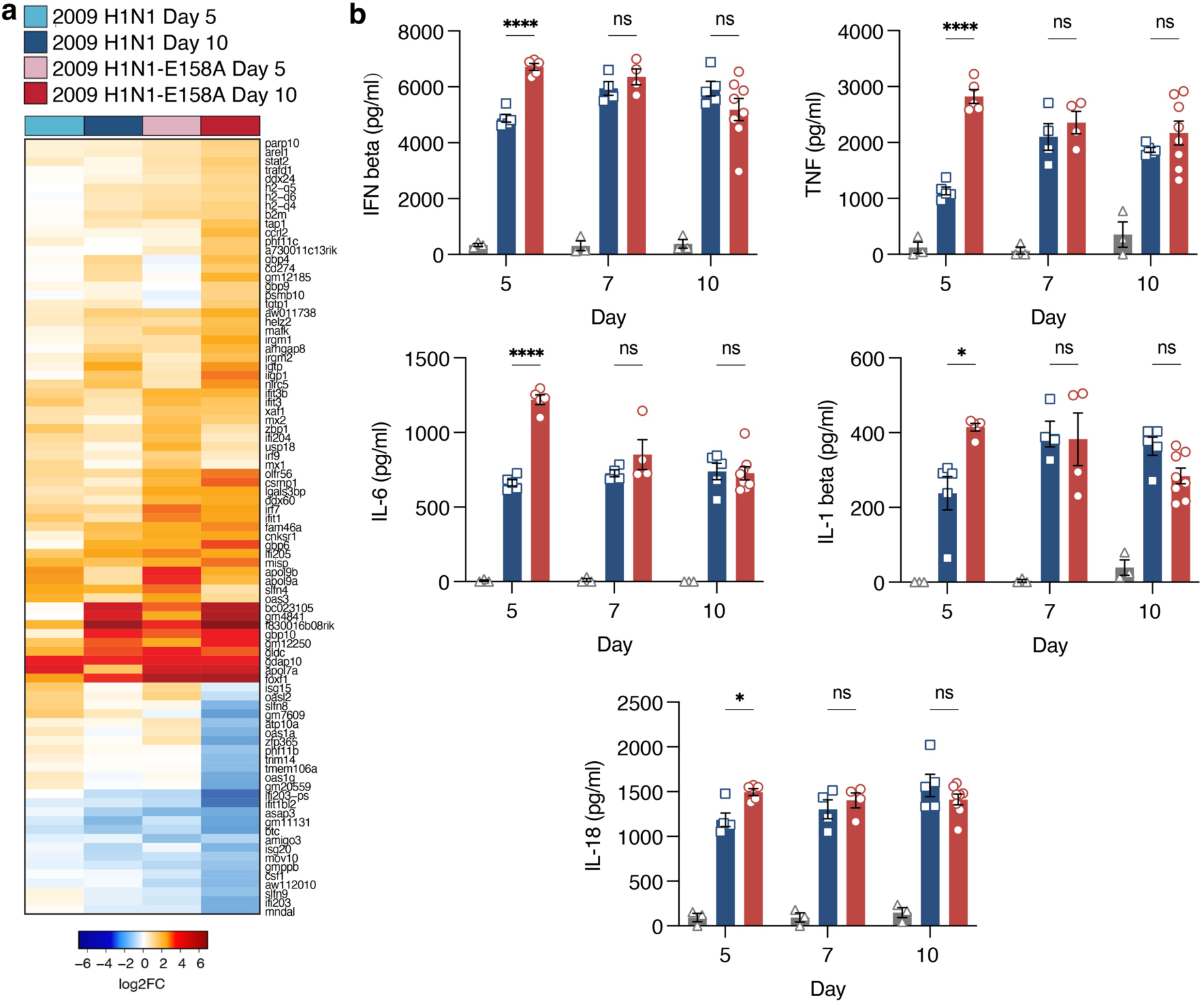
Mouse adapted 2009 H1N1 virus induces cardiac antiviral and inflammatory responses. **a,** Bulk RNA sequencing data from infected mice relative to mock controls (as in Fig. 4) were analyzed to generate a heat map of Heatmap of 87 interferon-stimulated genes (ISGs) identified from the union of differentially expressed genes across experimental groups. **b,** Quantification of IFN-β, TNF, IL-6, IL-1β, and IL-18 levels in heart homogenates from mice inoculated with PBS (Mock) or 200 TCID50 H1N1 virus, or H1N1-E158A virus, measured by ELISA at days 5, 7, and 10 post-infection. Each data point represents a single mouse; Error bars indicate SEM. Group sizes: Mock (days 5, 7, and 10), n = 3; H1N1: day 5, n = 5; day 7, n = 4; day 10, n = 5; H1N1-E158A: day 5, n = 5; day 7, n = 4; day 10, n = 8. Data are from a single experiment. Statistical significance was determined by two-way ANOVA with Tukey’s multiple comparisons test; *P < 0.05, ****P < 0.0001.

**Supp Fig 5:**
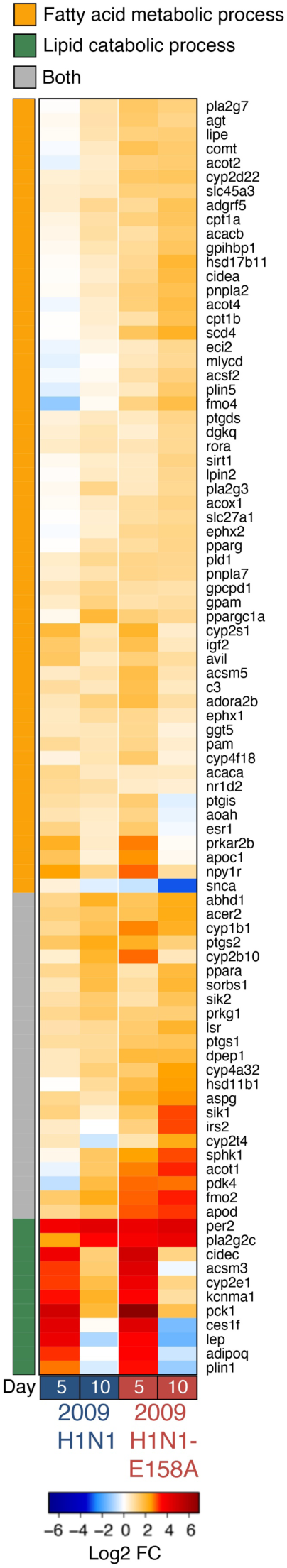
RNA sequencing reveals alteration of lipid metabolism-associated genes in influenza virus infected hearts. Heat map showing the relative expression of genes involved in cardiac lipid metabolism and mitochondrial pathways at days 5 and 10 post-infection across experimental groups.

**Supp Fig 6:**
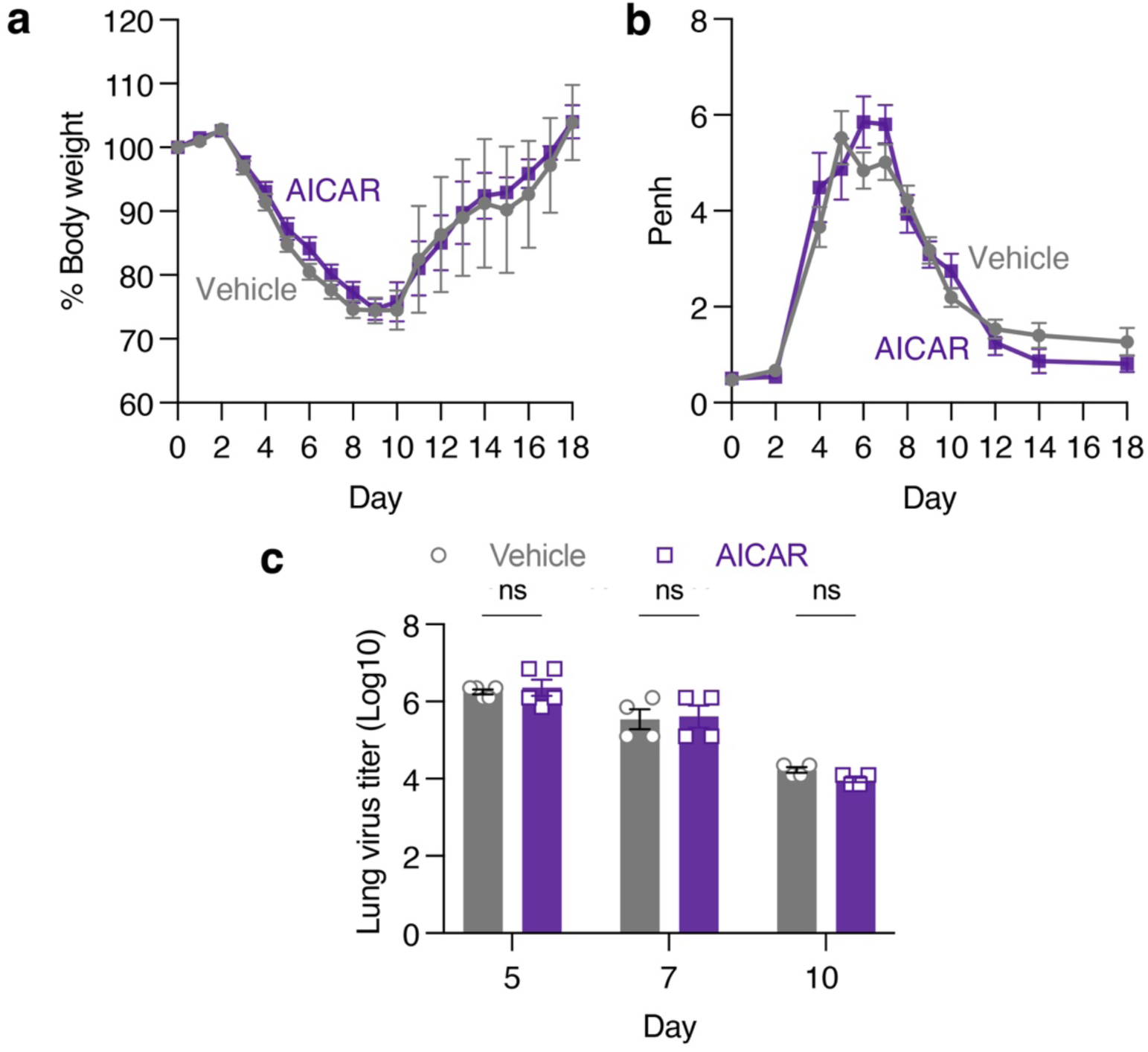
AICAR treatment does not affect weight loss, lung function, or lung viral titer in influenza virus infection. Mice infected with H1N1-E158A (200 TCID50) were treated daily with AICAR (100 mg/kg in saline) or vehicle (saline) from day 4 through day 10 post-infection. **a,** Body weight loss over time following infection. Each data point represents the average weight of individual mice normalized to 100% of baseline (day 0); error bars indicate SEM. Data from two experiments (Vehicle: n = 20; AICAR-treated: n = 16). **b,** Whole-body plethysmography measurements of PenH as a marker of respiratory function. Each data point represents the average Penh value from multiple mice; error bars indicate SEM. Data from two experiments (Vehicle: n = 20; AICAR-treated: n = 16). **c,** Lung viral titers at days 5, 7, and 10 post-infection in H1N1-E158A infected mice treated with AICAR or saline (vehicle). Data are from a single experiment. Group sizes: AICAR and vehicle groups, day 5, n = 5; days 7 and 10, n = 4. Statistical analysis was evaluated by two-way ANOVA followed by Sidak’s multiple comparisons test.

**Supp Fig 7:**
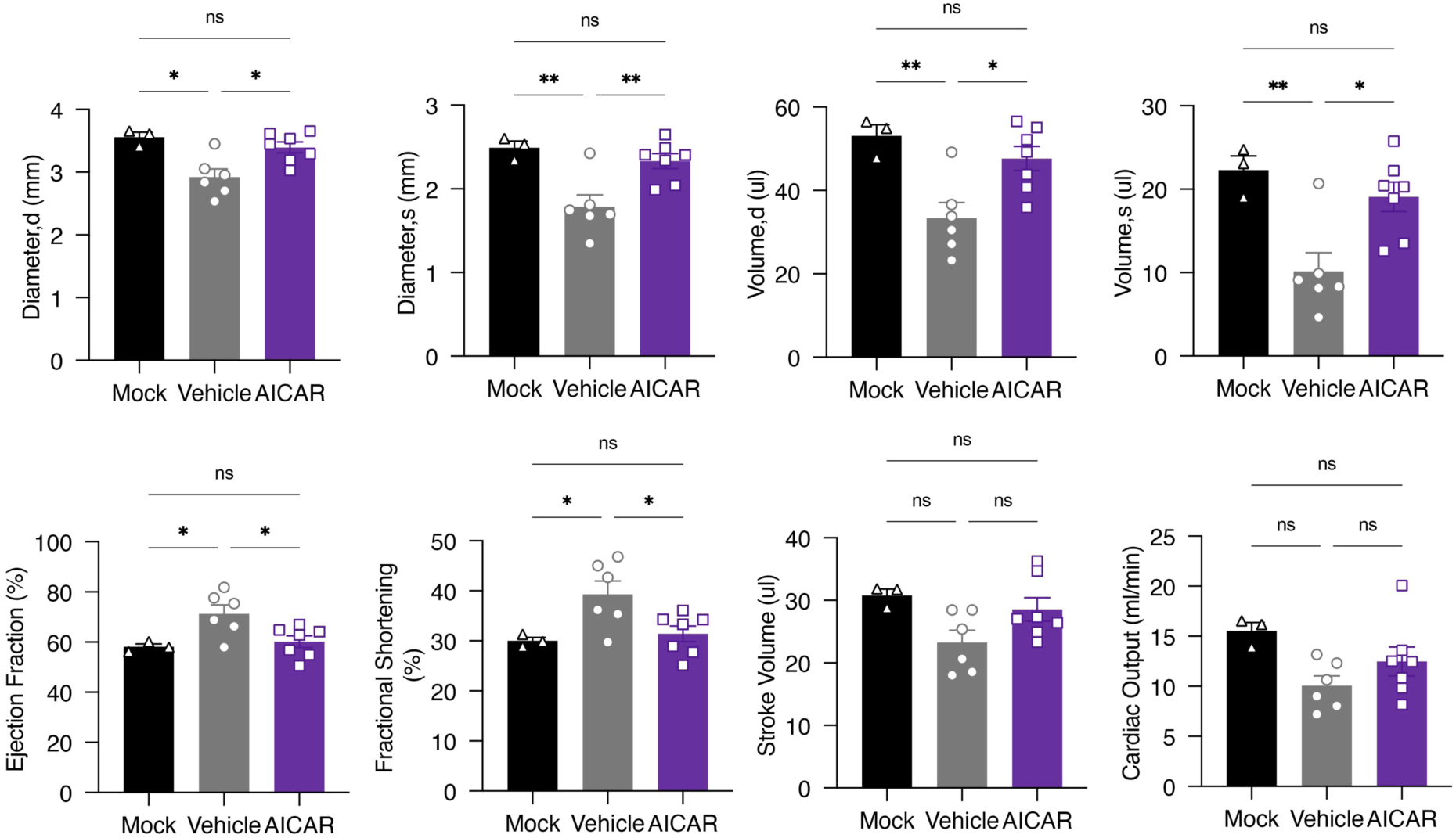
AICAR treatment normalizes key echocardiography parameters in influenza virus infected mice. Mice infected with H1N1-E158A (200 TCID50) were treated daily with AICAR (100 mg/kg in saline) or vehicle (saline) from day 4 through day 10 post-infection. Cardiac morphology and function were evaluated by 2-D echocardiography at 10 days post infection or on mock infected animals. Standard echocardiography readout data are plotted with each dot corresponding to an individual mouse. Error bars indicate SEM (Pre-infection: Mock n = 3, Vehicle n = 6, AICAR n = 7; Statistical significance was determined by one-way ANOVA with Tukey’s multiple comparisons test; *P < 0.05, **P < 0.01.

